# Midgut Transcriptome Analysis of *Clostera anachoreta* Treated with Cry1Ac Toxin

**DOI:** 10.1101/568337

**Authors:** Liu Jie, Wang Liucheng, Zhou Guona, Liu Junxia, Gao Baojia

## Abstract

*Clostera anachoreta* is one of the important Lepidoptera insect pests in forestry, especially in poplars woods in China, Europe, Japan and India, et al, and also the target insect of Cry1Ac toxin and Bt plants. In this study, by using the different dosages of Btcry1Ac toxin to feed larvae and analyzing the transcriptome data, we found six genes, HSC70, GNB2L/RACK1, PNLIP, BI1-like, arylphorin type 2 and PKM, might be associated with Cry1Ac toxin. And PI3K-Akt pathway, which was highly enriched in DEGs and linked to several crucial pathways, including the B cell receptor signaling pathway, toll-like receptor pathway, and MAPK signaling pathway, might be involved in the recovery stage when response to sub-lethal Cry1Ac toxin. This is the first study conducted to specifically investigate *C. anachoreta* response to Cry toxin stress using large-scale sequencing technologies, and the results highlighted some important genes and pathway that could be involved in Btcry1Ac resistance development or could serve as targets for biologically-based control mechanisms of this insect pest.

## 1. Introduction

Insect pest control in forestry and agriculture is increasingly achieved using biological, rather than chemical, insecticides, and the most successful of these is *Bacillus thuringiensis* (Bt). However, abuse of Bt pesticides and large-scale planting of Bt plants has caused enormous challenges and threats, including insect resistance and tolerance. Laboratory-selected resistant insect populations have shown that resistance can develop through multiple mechanisms, including alteration of Cry toxin activation (Oppert et al., 1997), toxin sequestration by lipophorin (Ma et al., 2005) or esterases (Gunning et al., 2005), elevation of immune response (Hernández-Martínez et al., 2010), and alteration of toxin receptors, resulting in reduced binding sites (Griffits and Aroian, 2005).

Exposure to high doses of Bt Cry toxin formulations, including feeding on highly expressing plants, can cause genetic mutations or produce higher fitness costs in target insects. Several reports have shown that after feeding on Bt toxin, evident fitness costs were associated with growth, development, and reproduction in resistant *Helicoverpa armigera*, *Heliothis virescens*, and *Plutella xylostella* strains, reducing spawning, hatch rate, feather rate, and larval survival indices, lengthening the development duration, and increasing larvae resistance levels. Aminopeptidase N (APN), cadherin-like protein (CAD), membrane-bound alkaline phosphatase (mALP), and ATP-binding cassette (ABC) transporter genes are the potential receptors of Bt Cry toxin, and their downregulated expression or mutation are the main cause of resistance. Previous reports have demonstrated that downregulation of the APN gene was related to Bt Cry1 resistance in *Spodoptera exigua* (Herrero et al., 2005; Park et al., 2014) and *Pectinophora gossypiella* (Fabrick et al., 2011), and reduced levels of mALP was a common feature in Cry-resistant strains of *P. xylostella*, *H. virescens*, *H. armigera*, and *Spodoptera frugiperda* when compared to susceptible larvae (Zhaojiang Guo, 2016; Jurat Fuentes and Asang, 2004; Jurat Fuentes et al., 2011). Mutation of cadherin and ABCC2 was genetically linked to Cry1Ac resistance and correlated with loss of Cry1Ac binding to membrane vesicles (Gahan et.al., 2001, Gahan et.al., 2010, Park et al., 2014). Two additional ABC transporter genes (PxABCG1 and ABCC1-3) in *P. xylostella* were trans-regulated by a constitutively transcriptionally activated upstream gene (PxMAP4K4) in the mitogen activated protein kinase (MAPK) signaling pathway outside the Bt R-1 resistance locus (Zhaojiang Guo, 2016.).

While prolonged exposure to sub-lethal doses of the toxin inhibits growth, development, and reproduction of larvae, imposing a fitness cost, it might also promote the tolerance and resistance of insects to toxins. An earlier report indicated that sub-lethal doses of toxin had deleterious effects on growth and recovery of *Achaea janata*; however, the insects exhibited epithelial cell proliferation in midgut tissue that coincided with elevated arylphorin expression (Chauhan VK, 2017), similar to the results reported by Castagnola in *H. virescens* (Anais Castagnola, 2017). These findings identified alpha-arylphorin as a protein with a putative role in response to Cry1Ac intoxication. Another report showed that serine proteases, detoxification enzymes (e.g., CarE, glutathione S-transferase (GST), and cytochrome P450 (P450)), and ABC transporters in *Cnaphalocrocis medinalis* midgut were differently expressed in response to Cry1C toxin. Antigen processing and expression, and the metabolic pathway of chronic myelogenous leukemia are highly involved in Bt response and metabolism (Yang Y, 2018).

In this study, the transcriptome data of *Clostera anachoreta* larvae treated with a continuous sub-lethal dose of Bt Cry1Ac toxin were used to identify Bt-related genes. The data were then combined with growth and development indices to analyze the regulation of related genes involved in digestion, metabolism, and fitness costs, to explore the key genes involved in the response to Cry1Ac treatment and to clarify the mechanisms of the response of target insects to Bt or Bt plants.

## 2. Methods

### 2.1. Insect rearing

*C. anachoreta* larvae were collected in wild poplar woods near a highway with no biological control in Baoding, Hebei, China, in May 2016. The larvae were continuously fed without toxins for five generations prior to experimentation. The moths were placed in paper boxes with nets and fed a 10% honey solution. Eggs laid on the mesh were removed and transferred to glass bottles containing mesh and detached leaves. The larvae that hatched from eggs were cultured on the detached leaves, and then used in the experiments. All insect cultures were kept at 26±1°C with 70–80% relative humidity (RH) and a natural photoperiod.

### 2.2. Treatment with Cry1Ac toxin

Third instar larvae were selected for the feeding experiments. The larvae were reared on poplar leaves dipped in different concentrations of Cry1Ac activated toxin solutions (Beijing Meiyan Agricultural Technology), resulting in growth inhibition and increased mortality of the larvae within 5 days. The larvae were treated with three different concentrations of Cry1Ac toxin: 15 μL/mL (Sample A), 7.5 μL/mL (Sample B), and 1.5 μL/mL (Sample C). Larvae treated with 5 Mm/L sodium carbonate solution were used as the negative control treatment. All larval treatments were incubated for 5 days at 26±1°C with 70–80% RH and a natural photoperiod, followed by feeding with control leaves under the same conditions until fourth instar for dissection. Each treatment group included three parallel samples, with approximately 10–15 larvae per bottle and 10–15 total bottles per treatment. Larvae were starved for 1 day prior to dissection. Dissected midguts were washed with diethyl pyrocarbonate water and blotted dry, then placed into centrifuge tubes (four to five midguts per tube), frozen with liquid nitrogen, and preserved at −80°C until use.

### 2.3. Sample collection and preparation

#### 2.3.1 RNA isolation, quantification and qualification

Total RNA was isolated from midgut of C. anachoreta larvae using TRIzol reagent (Invitro-gen,Carlsbad,CA,USA) and RNAprep pure Tissue kit (TIANGEN, DP431) following the manufacturer’s protocol. RNA degradation and contamination was monitored on 1% agarose gels. RNA purity was checked using the NanoPhotometer® spectrophotometer (IMPLEN, CA, USA). RNA concentration was measured using Qubit® RNA Assay Kit in Qubit® 2.0 Flurometer (Life Technologies, CA, USA). RNA integrity was assessed using the RNA Nano 6000 Assay Kit of the Agilent Bioanalyzer 2100 system (Agilent Technologies, CA, USA).

#### 2.3.2 Library preparation for Transcriptome sequencing

A total amount of 1.5 µg RNA per sample was used as input material for the RNA sample preparations. Sequencing libraries were generated using NEBNext® Ultra™ RNA Library Prep Kit for Illumina® (NEB, USA) following manufacturer’s recommendations and index codes were added to attribute sequences to each sample. Briefly, mRNA was purified from total RNA using poly-T oligo-attached magnetic beads. Fragmentation was carried out using divalent cations under elevated temperature in NEBNext First Strand Synthesis Reaction Buffer (5X). First strand cDNA was synthesized using random hexamer primer and M-MuLV Reverse Transcriptase (RNase H-). Second strand cDNA synthesis was subsequently performed using DNA Polymerase I and RNase H. Remaining overhangs were converted into blunt ends via exonuclease/polymerase activities. After adenylation of 3’ ends of DNA fragments, NEBNext Adaptor with hairpin loop structure were ligated to prepare for hybridization. In order to select cDNA fragments of preferentially 250-300 bp in length, the library fragments were purified with AMPure XP system (Beckman Coulter, Beverly, USA). Then 3 µl USER Enzyme (NEB, USA) was used with size-selected, adaptor-ligated cDNA at 37°C for 15 min followed by 5 min at 95 °C before PCR. Then PCR was performed with Phusion High-Fidelity DNA polymerase, Universal PCR primers and Index (X) Primer. At last, PCR products were purified (AMPure XP system) and library quality was assessed on the Agilent Bioanalyzer 2100 system.

#### 2.3.3 Clustering and sequencing (Novogene Experimental Department)

The clustering of the index-coded samples was performed on a cBot Cluster Generation System using TruSeq PE Cluster Kit v3-cBot-HS (Illumia) according to the manufacturer’s instructions. After cluster generation, the library preparations were sequenced on an Illumina Hiseq platform and paired-end reads were generated.

### 2.4. Data analysis

#### 2.4.1 Quality control

Raw data (raw reads) of fastq format were firstly processed through in-house perl scripts. In this step, clean data (clean reads) were obtained by removing reads containing adapter, reads containing ploy-N and low quality reads from raw data. At the same time, Q20, Q30, GC-content and sequence duplication level of the clean data were calculated. All the downstream analyses were based on clean data with high quality.

#### 2.4.2 Transcriptome assembly

The left files (read1 files) from all libraries/samples were pooled into one big left.fq file, and right files (read2 files) into one big right.fq file. Transcriptome assembly was accomplished based on the left.fq and right.fq using Trinity (Grabherr et al, 2011) with min_kmer_cov set to 2 by default and all other parameters set default.

#### 2.4.3 Gene functional annotation

Gene function was annotated based on the following databases: Nr (NCBI non-redundant protein sequences); Nt (NCBI non-redundant nucleotide sequences); Pfam (Protein family); KOG/COG (Clusters of Orthologous Groups of proteins); Swiss-Prot (A manually annotated and reviewed protein sequence database); KO (KEGG Ortholog database); GO (Gene Ontology).

#### 2.4.4 Quantification of gene expression levels

Gene expression levels were estimated by RSEM (Li et al, 2011) for each sample: 1. Clean data were mapped back onto the assembled transcriptome. 2. Readcount for each gene was obtained from the mapping results.

#### 2.4.5 Differential expression analysis

##### For the samples with biological replicates

Differential expression analysis of two conditions/groups was performed using the DESeq R package (1.10.1). DESeq provide statistical routines for determining differential expression in digital gene expression data using a model based on the negative binomial distribution. The resulting P values were adjusted using the Benjamini and Hochberg’s approach for controlling the false discovery rate. Genes with an adjusted P-value <0.05 found by DESeq were assigned as differentially expressed.

##### For the samples without biological replicates

Prior to differential gene expression analysis, for each sequenced library, the read counts were adjusted by edger (Robinson MD, 2010) program package through one scaling normalized factor. Differential expression analysis of two samples was performed using the DEGseq (Anders S, 2010) R package. P value was adjusted using q value (Storey et al, 2003). Q value<0.005 & |log2(foldchange)|>1 was set as the threshold for significantly differential expression.

#### 2.4.6 GO enrichment analysis

Gene Ontology (GO) enrichment analysis of the differentially expressed genes (DEGs) was implemented by the GOseq R packages based Wallenius non-central hyper-geometric distribution (Young et al, 2010), which can adjust for gene length bias in DEGs.

#### 2.4.7 KEGG pathway enrichment analysis

KEGG (Kanehisa et al., 2008) is a database resource for understanding high-level functions and utilities of the biological system, such as the cell, the organism and the ecosystem, from molecular-level information, especially large-scale molecular datasets generated by genome sequencing and other high-throughput experimental technologies (http://www.genome.jp/kegg/). We used KOBAS (Mao et al., 2005) software to test the statistical enrichment of differential expression genes in KEGG pathways.

## 3 Results

### 3.1 Analysis of growth and development indices

Continuous bioassays were conducted using *C. anachoreta* larvae in the same growth stage. Larval mortality and the developmental duration of third generation larvae were significantly affected by increased dosages of Cry1Ac. Insect and pupa weights were highest at 7.5 μg/mL(Sample B), and were almost consistent with the control weights (Figure 1).

**Figure 1.**
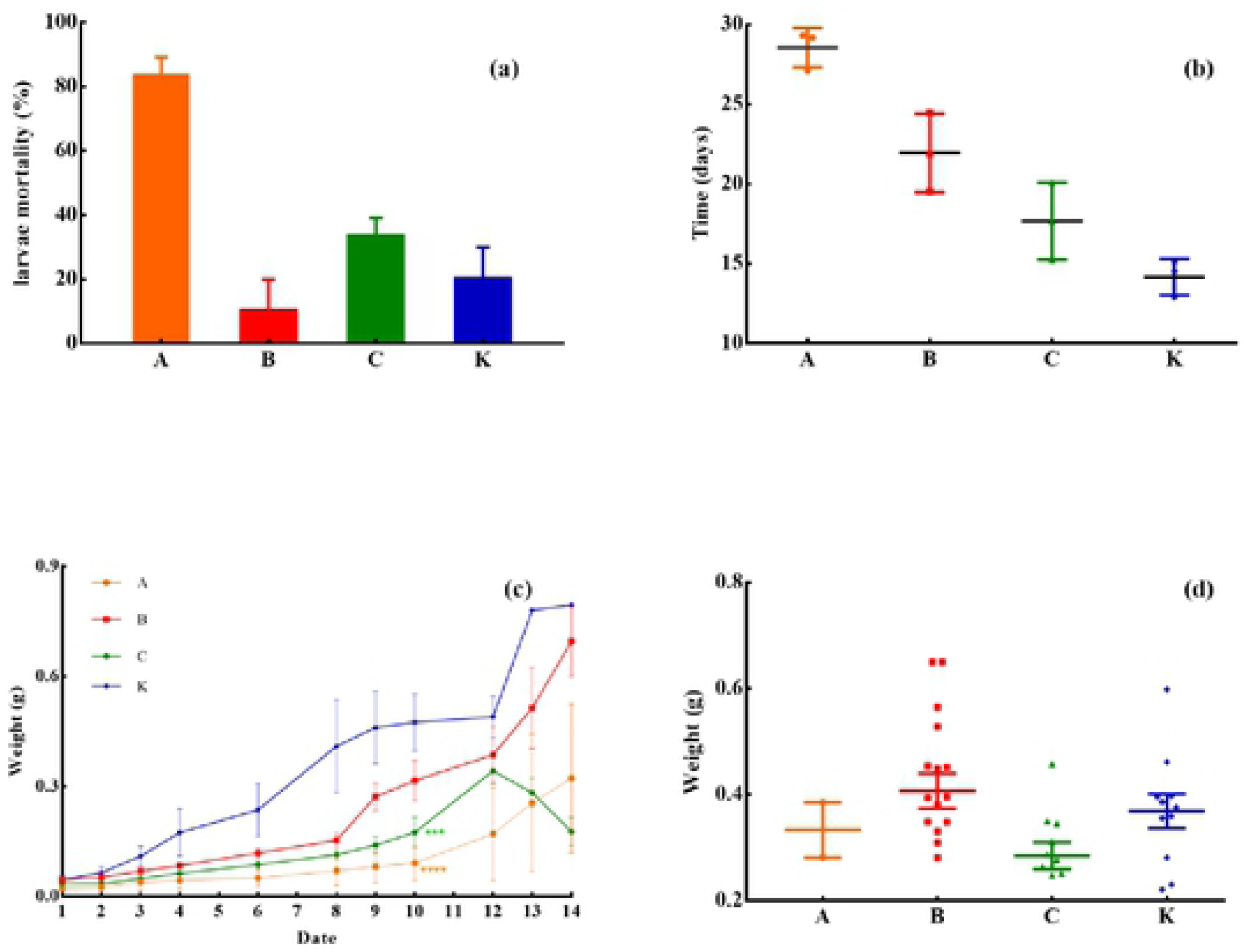
Growth and development indices of *C. anachoreta* larvae in response to Cry1Ac toxin.

### 3.2 Illumina Sequencing and de Novo Assembly

After sequencing using Illumina HiSeqTM 2000 system, we generated 57516756, 56506756, 45779646 and 46263752 raw reads in Sample A, B, C and K (Control) respectively. After removal of low-quality reads and reads containing adaptor or N (It means the bases in reads were not certain), the libraries yielded totals of 54563000, 53586232, 43332730 and 44893556 clean reads. Their Q30 values (meaning that the base recognition accuracy is 99.9%) were 89.25%, 91.51%, 89.07% and 91.67%, respectively (Table 1). Trinity software (Grabherr M.G.et al., 2013) was used to assemble the clean reads, producing 219275 assembled sequences (transcripts) with a total length of 213501751, an average length of 974 and an N50 of 1889. After clustered by TGICL (Pertea G. et al., 2003), a total of 151,090 unigenes with a total length of 193414771, an average length of 1280 and N50 of 2125 were obtained (Table 1). The sequencing raw data was not uploaded.

**Table 1.** Summary of the *C. anachoreta* midgut transcriptome

| Description | A | B | C | K | Both |
| --- | --- | --- | --- | --- | --- |
| RIN (RNA integrity number) | 9.7 | 9.3 | 9.1 | 9.6 |  |
| Raw Reads | 57516756 | 56506756 | 45779646 | 46263752 |  |
| Clean reads | 54563000 | 53586232 | 43332730 | 44893556 |  |
| Clean bases | 8.18G | 8.04G | 6.5G | 6.73G |  |
| Q30 (%) | 89.25 | 91.51 | 89.07 | 91.67 |  |
| Total number of unigenes |  |  |  |  | 151090 |
| Total length of unigenes |  |  |  |  | 193,414,771 |
| Max length of unigenes |  |  |  |  | 29377 |
| Min length of unigenes |  |  |  |  | 201 |
| Mean length of unigenes |  |  |  |  | 1280 |
| N50 |  |  |  |  | 2125 |

All assembled unigenes were more than 200 bp in size and the maximum length of unigenes was 29377 bp. Among these unigenes, 89,559 (59.28%) were in the length range of 200 to 1000, 32407 (21.45%) were in the length range of 1000 to 2000 and 29,124 (19.28%) were longer than 2000 bp (Figure 2). All of the data indicated that the integrity of unigenes was relatively good.

**Figure 2.**
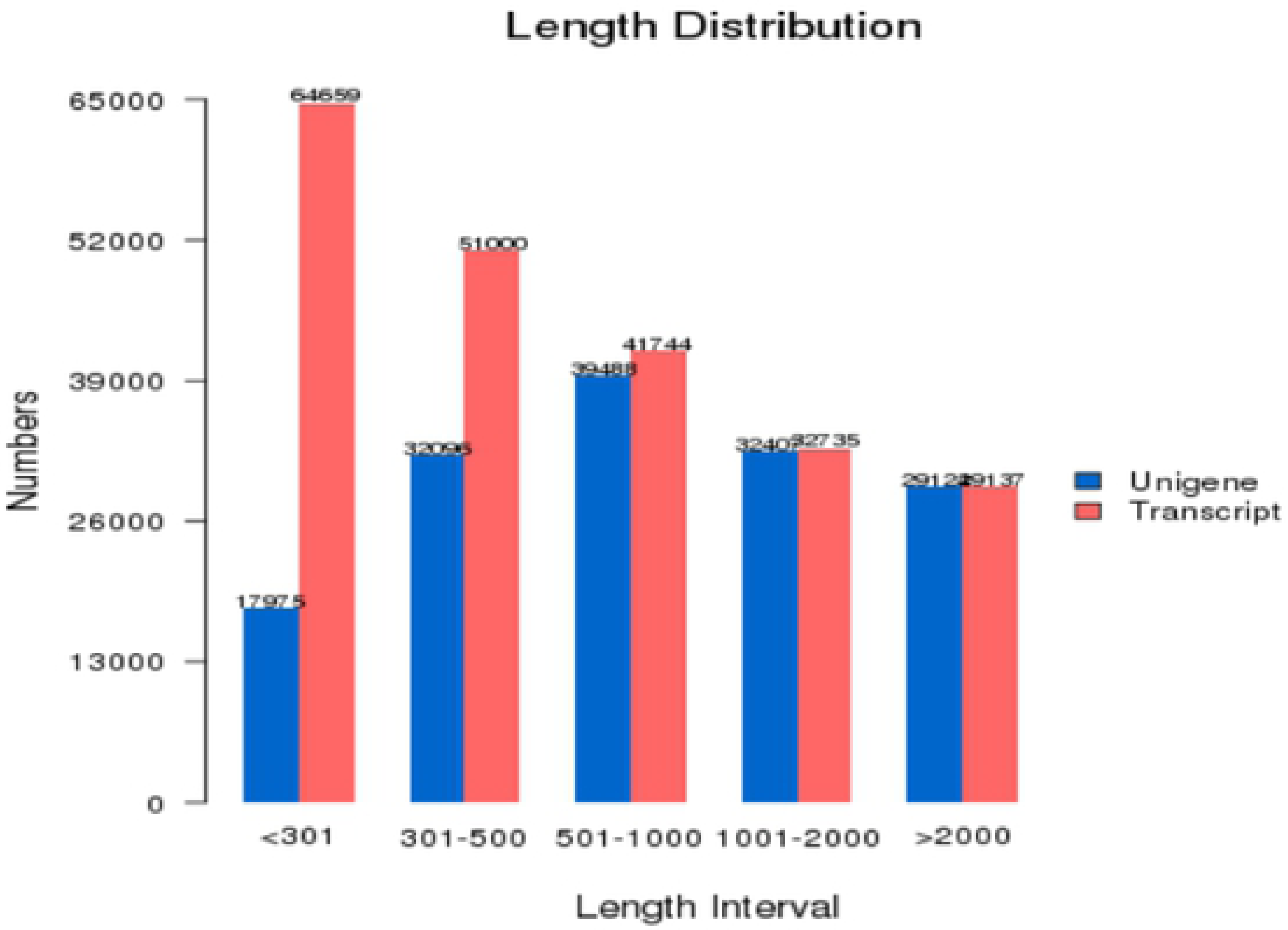
Histogram of the unigenes and transcrpts length distribution

### 3.3 Assembly, evaluation, and annotation

Of 90,682 unigenes, 41.96%, 32.8%, 16.21%, 31.13%, 35.74%, 36.14%, and 19.08% received significant hits in the non-redundant (Nr), nucleotide (NT), Kyoto Encyclopedia of Genes and Genomes (KEGG) Orthology (KO), Swiss-Prot, Pfam, Gene Ontology (GO), and Eukaryotic Clusters of Orthologous Groups (KOG) databases, respectively (Table 2). Using the online software Novomagic (Novogene), the numbers of unigenes that were unique or shared among the databases were visualized with a Venn diagram (Figure 3). Among them, 10,946 unigenes had reference sequences in all seven databases, while 90,682 (60.01%) unigenes were aligned to at least one database. However, 60,408 (39.99%) unigenes could not be annotated, likely because they included a large number of novel genes or non-coding RNA sequences. These non-annotated unigenes might also have been too short to generate sequence matches. A total of 63,400 unigenes were aligned to the Nr database; the E-value distribution of these unigenes showed that 5.1% had perfect matches, while 52.0% showed homologies ranging from 1×10^−45^ to 1×10^−5^. In addition, the species distribution indicated that ≥ 64.1% of the unigenes matched sequences from Lepidoptera insects; 36.1% had the highest homology to *Bombyx mori*, followed by *Danaus plexippus* (13.7%), *P. xylostella* (12.6%), and *Ostrinia nubilalis* (1.7%) (Figure 4). This was consistent with the genetic relationships among these species, supporting the validity of our transcriptome data.

**Figure 3.**
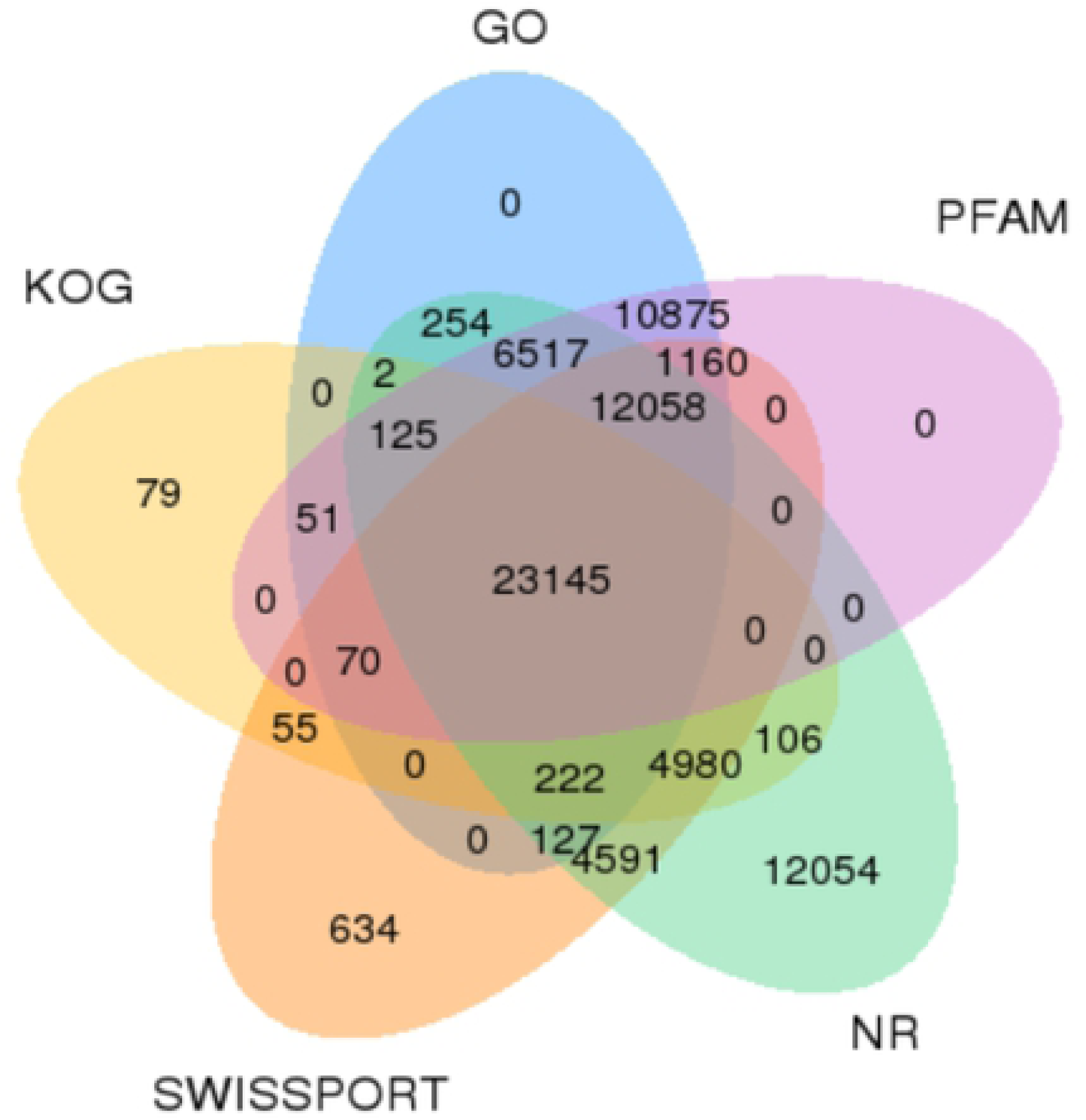
Venn diagram showing the numbers of unique and sharedannotated unigenes among five public databases.

**Figure 4.**
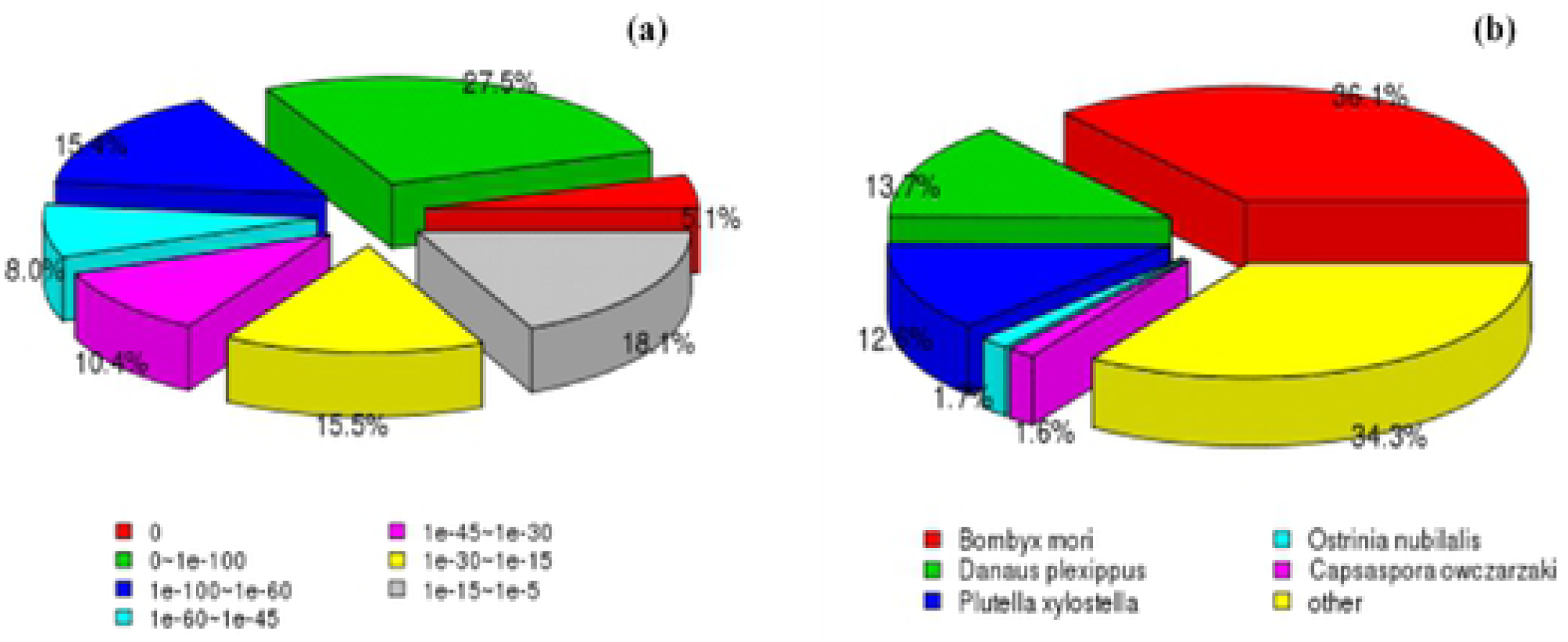
E-value distribution and species distribution of assembled unigenes aligned to the Nr database. (a) E-value distribution; (b) Species classification

**Table 2.** Annotation of unigenes BLAST-queried against seven public databases

|  | Number of Genes | Percentage (%) |
| --- | --- | --- |
| Annotated in NR | 63400 | 41.96 |
| Annotated in NT | 49563 | 32.8 |
| Annotated in KO | 24500 | 16.21 |
| Annotated in Swiss-Prot | 47042 | 31.13 |
| Annotated in PFAM | 54001 | 35.74 |
| Annotated in GO | 54606 | 36.14 |
| Annotated in KOG | 28835 | 19.08 |
| Total Unigenes | 151090 | 100 |

### 3.4 KOG Analysis

Using the KOG databases, we predicted the function of the unigenes and classified them according to possible functions. Among the 25 KOG categories, the cluster for “General function prediction only” represented the largest group (4517, 15.66%), followed by “Signal transduction mechanisms” (3431, 11.90%) and “Posttranslational modification, protein turnover, chaperones” (3071, 10.65%). Expect the “X, Unamed protein” obtained 2 unigenes that we do not know the function was, the smallest groups were “Cell motility” (45, 0.16%) and “Nuclear structure” (164, 0.57%).

Figure 5 also showed that 205 and 320 unigenes were involved in the cluster for “Defense mechanisms” and “Cell wall/membrane/envelope biogenesis”, respectively. And 5304 unigenes were involved in the cluster for Digestion and absorption consisted of “Energy production and conversion”, “Lipid transport and metabolism”, “Carbohydrate transport and metabolism” and “Amino acid transport and metabolism”. In this experiment, the largest group was “General function prediction only” which the same as many other studies (Tang, C. et al.,2013; Wang, X.W,et al., 2010; Xu, Q. et.al., 2013), and the follow group was “signal transduction mechanisms” which might be more important for *C. anachoreta* responding to BtCry1Ac in midgut. With this in mind, we paid more attention to the signal transduction mechanisms. Furthermore, many of the unigenes participated in defense mechanisms and digestion and absorption, which might imply that the organism would need much more energy to defend foreign substances after dieting Cry1Ac toxin.

**Figure 5.**
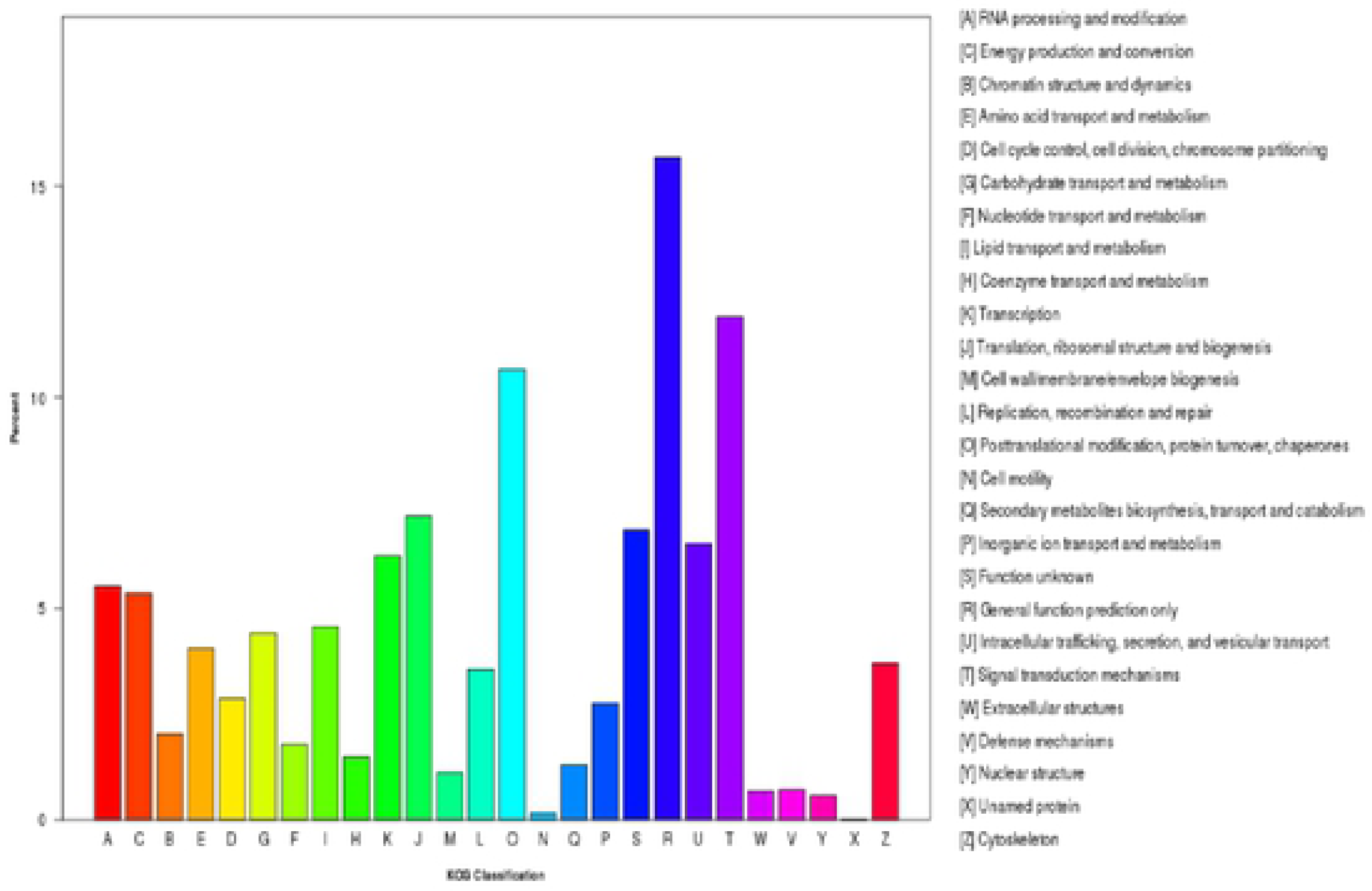
Histogram presentation of eukaryotic clusters of orthologous groups (KOG) classifications.

### 3.5 GO Analysis

Using Blast2GO v2.5 software (Götz S, 2008), we classified these sequences at two levels. Finally, 54606 transcripts were divided into three main categories (CC cellular component, MF molecular function, BP biological process) and 54 subcategories based on my data. In the subcategories of CC, most of the corresponding genes were involved in “cell” (7563, GO:0005623), “cell part” (10433, GO:0044464) and “organelle” (7179, GO:0043226) (Figure 6). In the categories of MF, the largest term is “binding” (GO:0005488) into which 11881 unigenes were classified, followed by the terms “catalytic activity” (15647, GO:0003824). In the BP category, we observed that 201 and 4936 unigenes belonged to the categories “immune system process” and “response to stimulus”, respectively, 20493 unigenes were participated in the “metabolic process” (GO:0008152). It was noteworthy that in GO analysis, one unigene could be classified into more than one GO term due to its multiple functions. Hence, there were common unigenes in these three catergories “immune system process”, “response to stimulus” and “metabolic process” (Figure 7). Nevertheless, 2913 unigenes (14.21%) in the cluster of “metabolic process” were related to the stress or defense response, only 0.04% and 3.5% unigenes were divided into in the cluster of “immune system process” and “response to stimulus”, respectively (Tariq, M. et al., 2015) The midgut chosen for our study contained the certain receptor protein of Cry1Ac, which was dissected and extracted from larvae dieting with Cry1Ac toxin in different doses, might be an important site of resistance or defense response to Cry1Ac. It could explain the difference of the expression between each sample involved in resistance or defense system (Figure 8).

**Figure 6.**
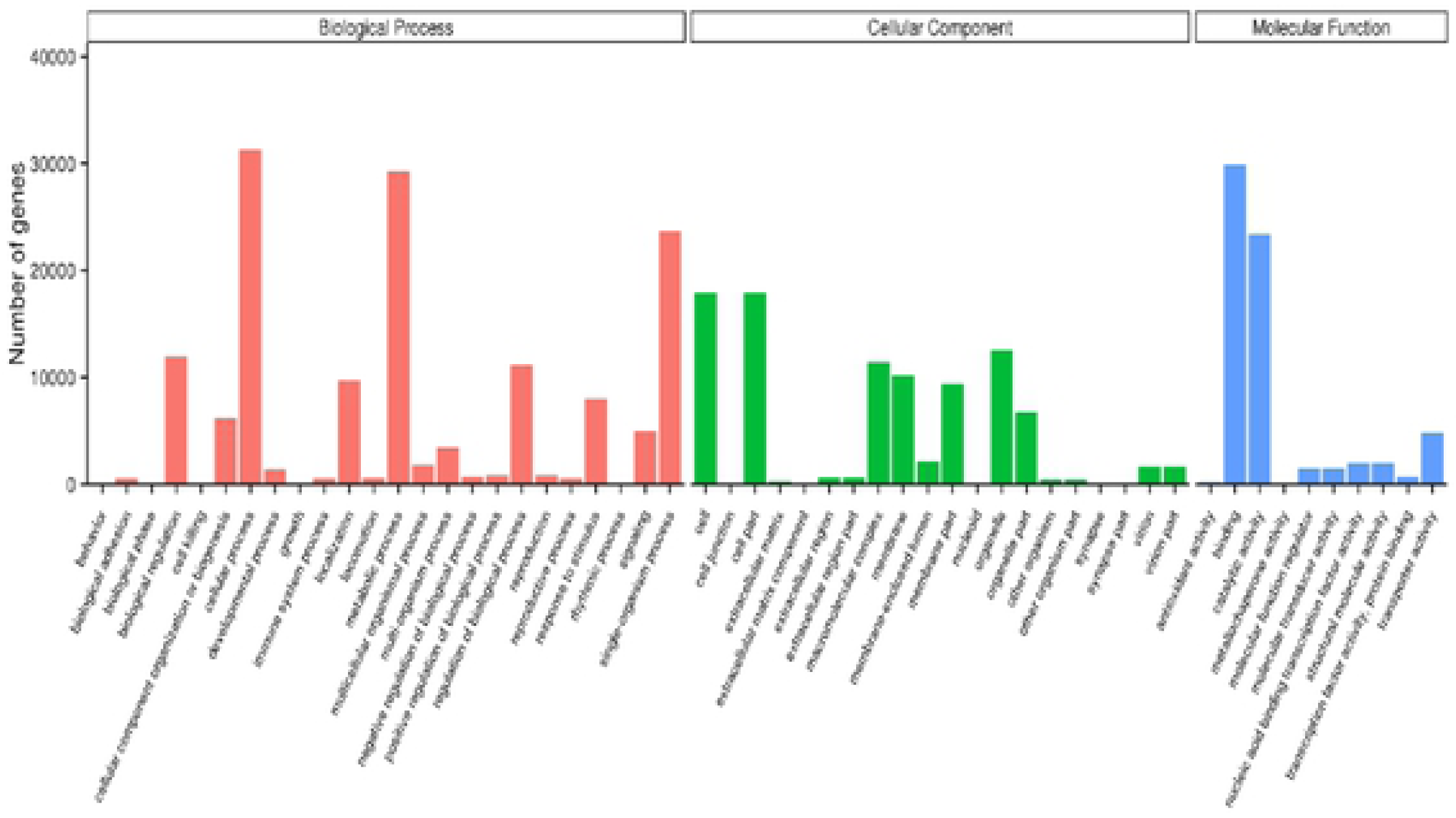
Histogram presentation of Gene Ontology (GO) classifications.

**Figure 7.**
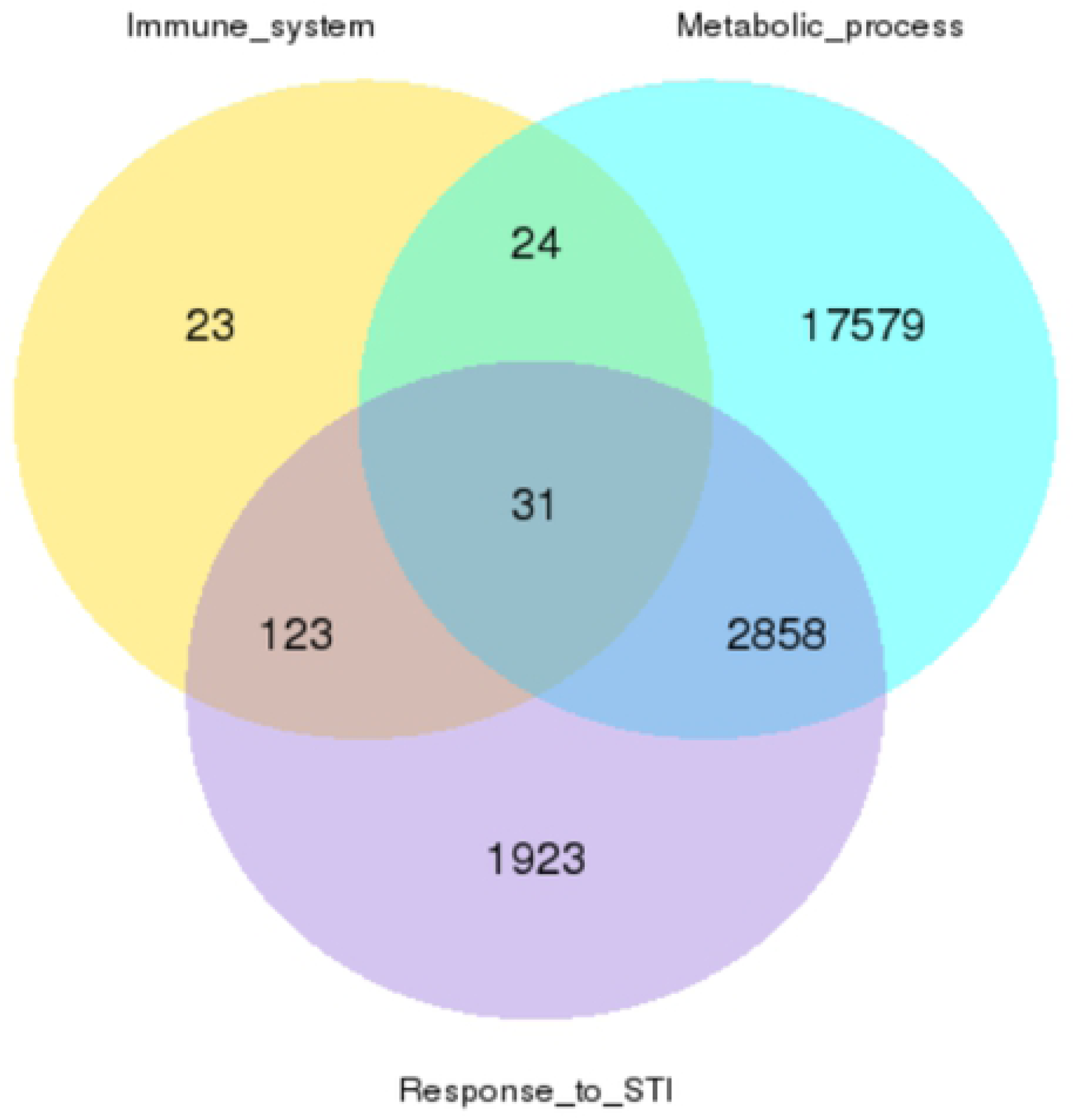
Shared and unique unigenes in three GO categories: Immune system, Metabolic Process, and Response to stimulus.

**Figure 8.**
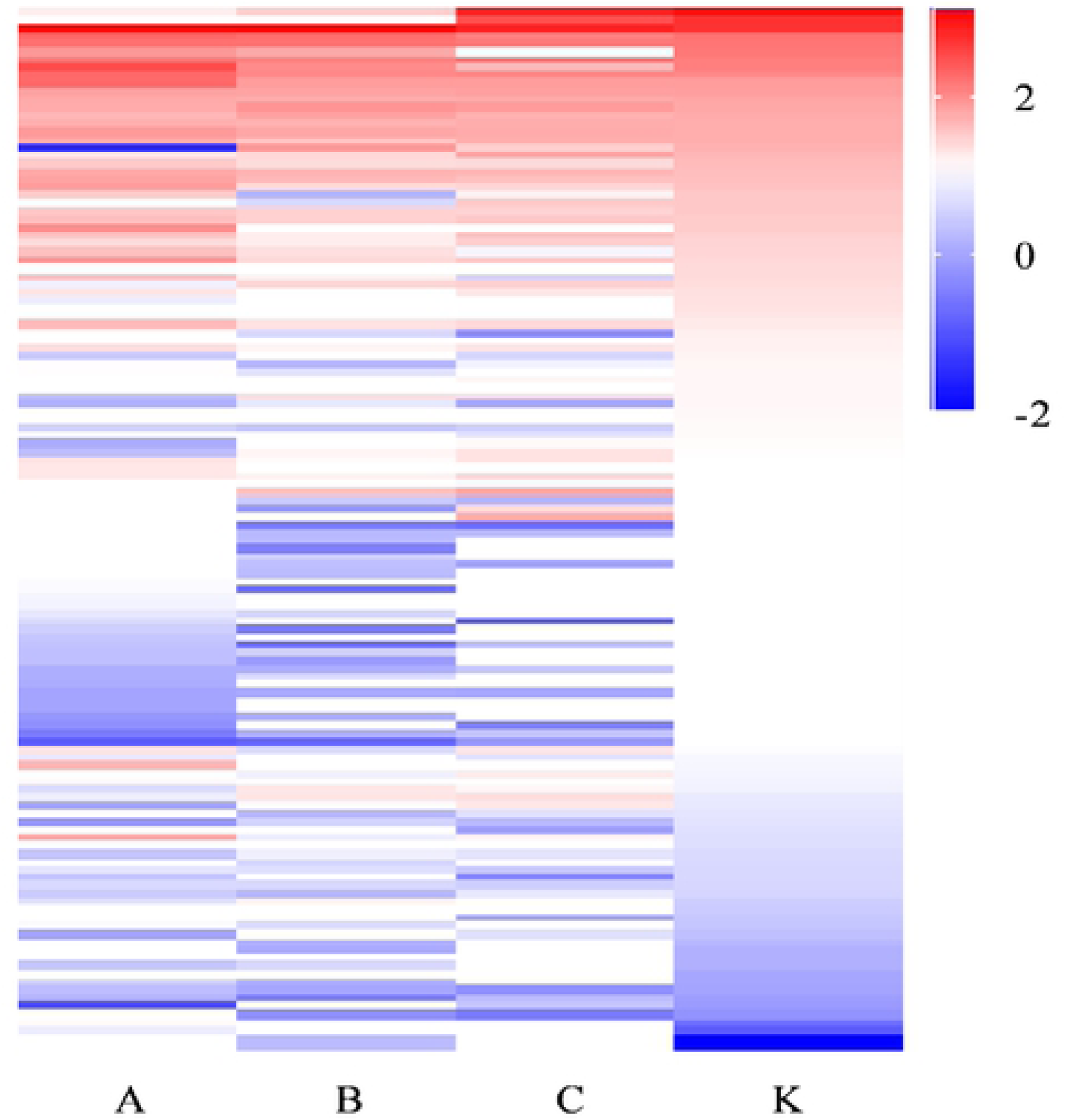
Heatmap of FPKM values for unigenes related to the immune system.

### 3.6 KEGG analysis

Unigenes were queried against the KEGG database to identify the biological processes in which they are involved, and a total of 14,829 unigenes were assigned to 231 KEGG pathways. For this study, 5998 unigenes were assigned to 45 KEGG pathways of interest. The top three represented pathways in the “Environmental Information Processing” cluster were the “PI3K-Akt signaling pathway” (478 unigenes), “cAMP signaling pathway” (468 unigenes), and “MAPK signaling pathway” (459 unigenes). The top three pathways in the “Metabolism” cluster were “Oxidative phosphorylation,” “Glycolysis/Gluconeogenesis,” and “Galactose metabolism,” all of which are related to digestion and absorption. The pathways related to resistance or tolerance of Cry1Ac toxin were “Metabolism of xenobiotics by cytochrome P450,” “Drug metabolism-other enzymes and P450,” and “Degradation of aromatic compounds,” with 165, 287, and 42 unigenes, respectively. In the “Cell growth and death” category, the “p53 signaling pathway” had 119 unigenes. The “Ubiquitin mediated proteolysis” pathway had 357 unigenes. There were seven pathways represented in the “Immune system” category: “Chemokine signaling pathway” (273 unigenes), “Leukocyte transendothelial migration” (240 unigenes), “T cell receptor signaling pathway” (210 unigenes), “Antigen processing and presentation” (179 unigenes), “Toll-like receptor signaling pathway” (108 unigenes), “B cell receptor signaling pathway” (91 unigenes), and “Intestinal immune network for IgA production” (2 unigenes) (Figure 9).

**Figure 9.**
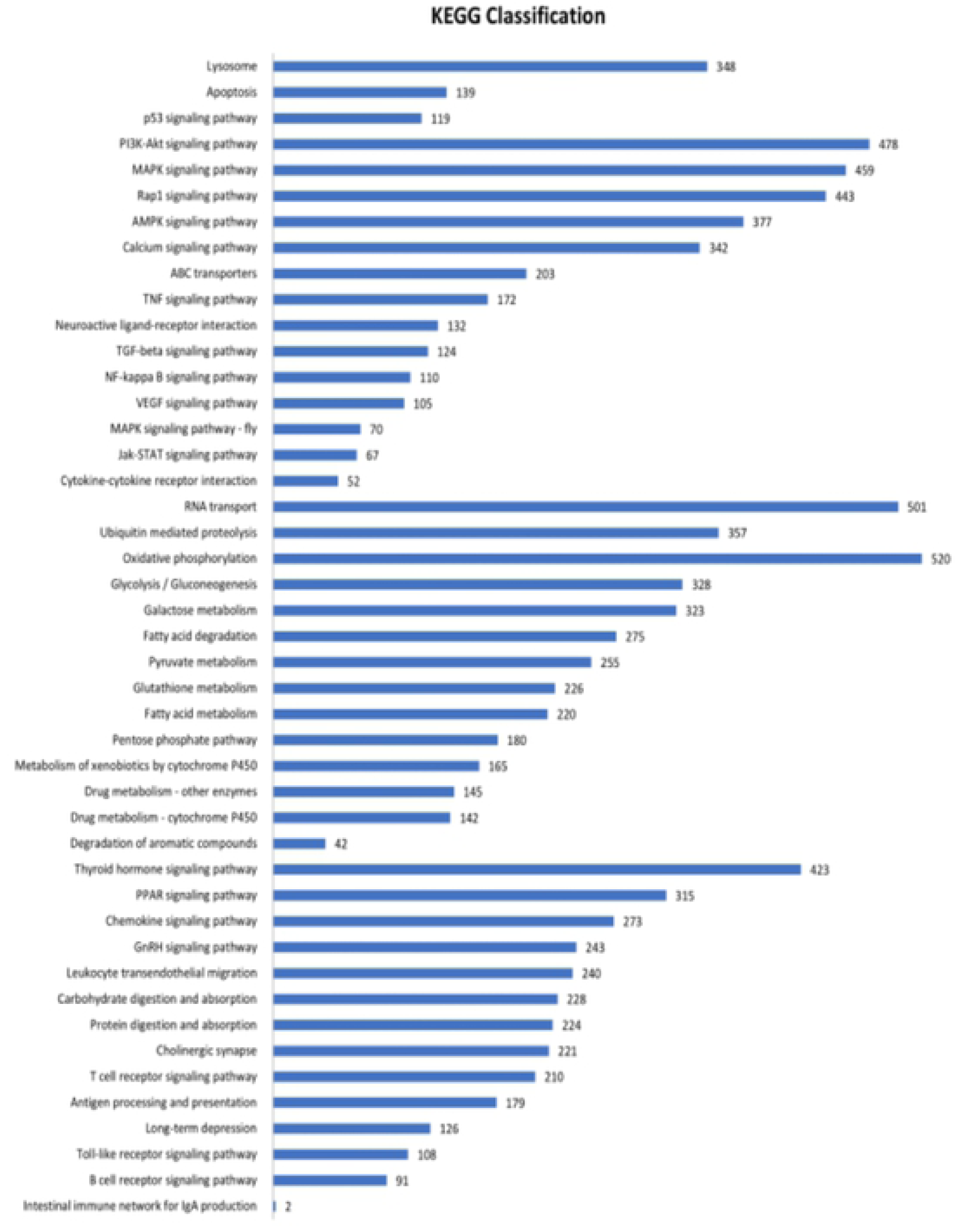
KEGG classifications related to Cry1Ac toxin.

### 3.7 Analysis of differentially expressed genes (DEGs)

In the midgut of *C. anachoreta* treated with Cry1Ac, the three treatments (A, B, and C) compared to the control (K) had 1119, 615, and 261 upregulated unigenes, and 541, 560, and 301 downregulated unigenes, respectively (Figure 10). There were 158 DEGs in common among all three treatments, including 99 downregulated, 21 upregulated, and 38 unigenes with inconsistent expression (Supplementary Table 3) (Figures 11 & 12). Of the shared DEGs, six downregulated and three upregulated DEGs had explicit annotations, including Heat shock cognate protein 70 (HSC70), downregulated −9.69 to −5.51 fold; GNB2L/RACK1 (which binds to protein kinase C (PKC)), downregulated −8.24 to −1.95 fold; PNLIP (pancreatic lipase), downregulated −6.78 to −4.46 fold; Bax inhibitor-1-like protein (a BI1 family member and possible suppressor of apoptosis), upregulated 8.42 to 8.78 fold; and arylphorin type 2 (related to proliferation of epithelial cells), upregulated 2.97 to 7.34 fold (Table 4).

**Table 4.** Shared DEGs in midguts of *C. anachoreta* larvae treated with Cry1Ac toxin

| Gene_ID | Nr description (Protein name) | Species | A vs. K | B vs. K | C vs. K |
| --- | --- | --- | --- | --- | --- |
| Cluster-38636.56529 | HSC70 | *Trichoplusia ni* | -9.69 | -5.51 | -9.02 |
| Cluster-38636.2126 | GNB2L/RACK1 | *Bombyx mori* | -8.24 | -1.95 | -4.21 |
| Cluster-38636.102879 | myosin heavy chain 4 | *Danio rerio* | -6.81 | -7.06 | -6.84 |
| Cluster-38636.1660 | PNLIP | *Plutella xylostella* | -6.78 | -6.65 | -4.46 |
| Cluster-38636.34145 | larval cuticle protein 16/17-like | *Plutella xylostella* | -5.3 | -5.17 | -5.33 |
| Cluster-38636.53239 | epidermal growth factor receptor kinase substrate 8-like | *Bombyx mori* | -3.39 | -1.43 | -1.77 |
| Cluster-38636.56439 | Bax inhibitor-1-like protein | *Bombyx mori* | 8.42 | 8.63 | 8.78 |
| Cluster-38636.58350 | arylphorin type 2 | *Cerura vinula* | 5.11 | 7.34 | 2.97 |
| Cluster-38636.60574 | PKM, pyruvate kinase | *Papilio xuthus* | 4.81 | 6.31 | 6.58 |

Based GO enrichment analysis, most DEGs were involved in “Catalytic activity” and “Hydrolytic activity” under the “Molecular Function” category, except for Sample C versus the control; there was no significant difference between the control and the low-dose Cry1Ac treatment (Sample C).

### 3.8 Genes of interest and Bt-related genes in the transcriptome of *C. anachoreta*

We investigated DEGs potentially involved in insecticidal tolerance in *C. anachoreta*, including metabolism and Bt-related genes. DEG sequences encoding enzymes functioning in ingestion and detoxification as well as Cry1Ac receptors were extracted from the midgut RNA-Seq data. Cadherin, ABC transporter, ALP, aminopeptidase, P450, GST, carboxylesterase, and serine proteases including trypsin and chymotrypsin (FDR < 0.05, absolute value of log2FC ≥ 2) were significantly differentially expressed (Table 5). Arylphorin subunit beta and arylphorin type 2, related to regeneration of the midgut, also differed significantly between the Cry1Ac treatments and the control.

**Table 5.**
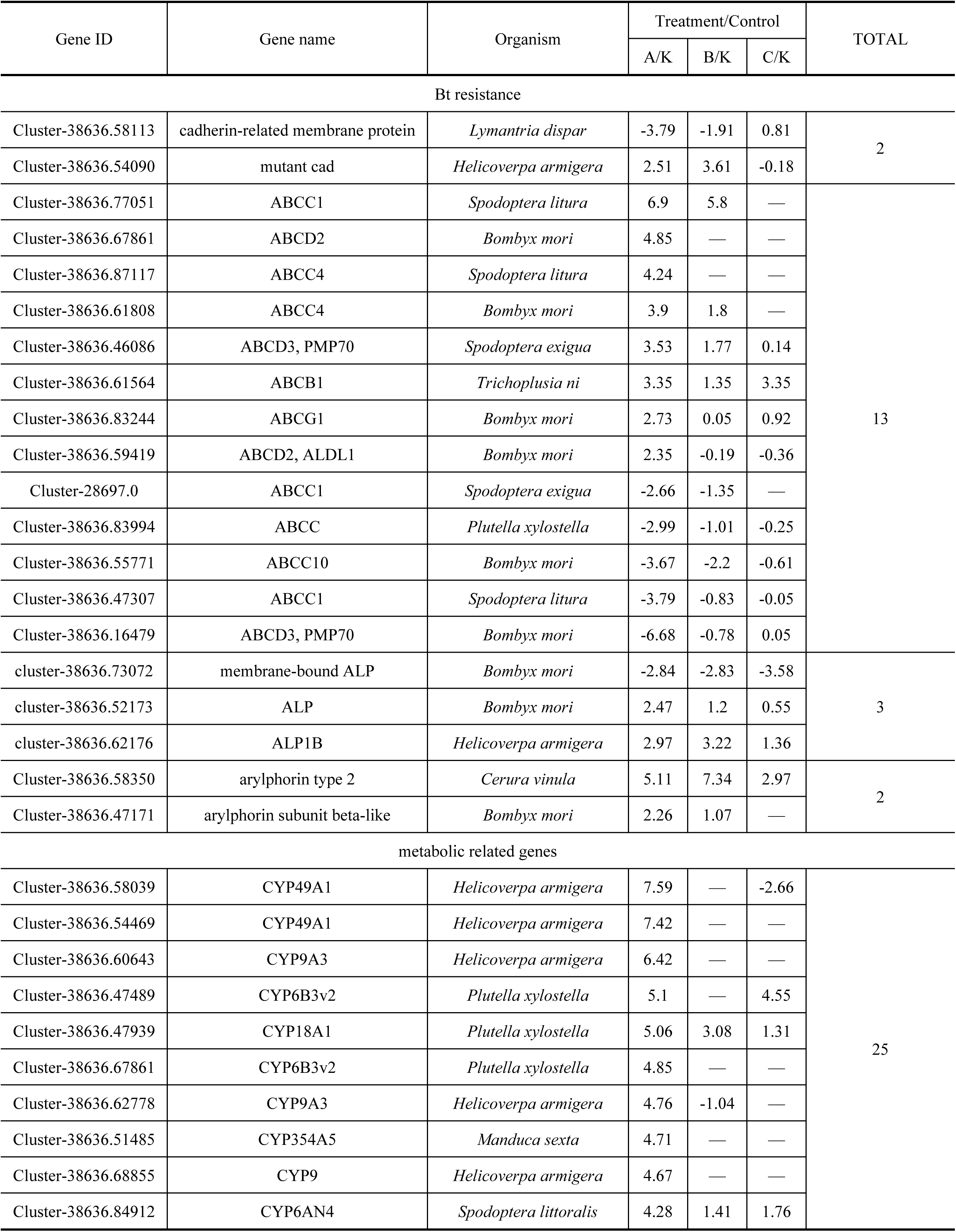
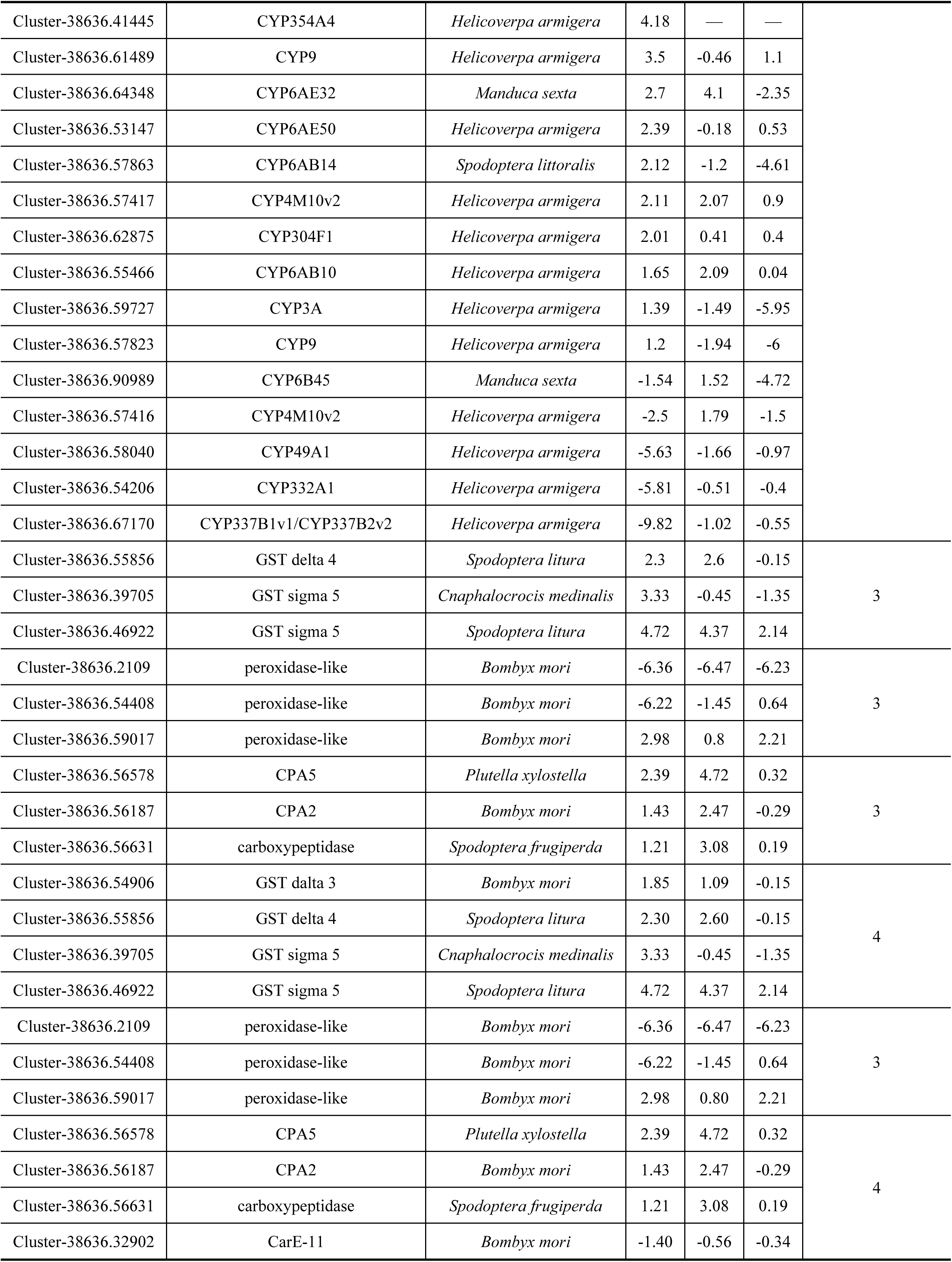
Differently expressed Bt-related genes potentially involved in the response of *C. anachoreta* larvae to Cry1Ac toxin

Of a total of 25 unigenes encoding CAD, only two were significantly differentially expressed: cadherin-related membrane protein was downregulated, while mutant CAD was upregulated in treatments A (2.51 fold) and B (3.61 fold). Of 47 total ABC transporter genes, 13 were differentially expressed in treatment A, while only one, Cluster-38636.61564 belonging to the ABCB1 family, was upregulated in all three treatments. Seven unigenes belonged to the ABCC family, of which four were downregulated in Samples A and B (−1.01 to −3.79 fold), while three were upregulated. Three unigenes were related to ALP, a potential receptor of Cry1Ac. Of them one gene, mALP, was downregulated in all three Cry1Ac treatments, while the others were upregulated or not significantly differentially expressed. Seven unigenes mapped to the APN family, a known receptor of Cry1Ac; however, they were not significantly differentially expressed. Two genes encoding arylphorin, related to cell regeneration, were significantly upregulated in Samples A and B (7.34 to 1.07 fold).

There were many unigenes involved in metabolic systems related to resistance/tolerance or immunity, including GST, P450, CarE, and serine proteases (Table 5). Of 238 total unigenes encoding P450, 25 were significantly differentially expressed in Sample A. Of these, 20 genes were upregulated in Sample A (from 1.20 to 7.59 fold), while five genes were downregulated (CYP337B1v1, CYP332A1, CYP49A1, CYP4M10v2, and CYP6B45). Notably, six genes, CYP49A1, CYP9A3, CYP6B3v2, CYP354A5, CYP9, and CYP354A4, were only expressed in Sample A. Of three GST genes, all were upregulated in Sample A, two in B, and one in C. Three CarE genes were significantly upregulated in Samples A and B, and three genes encoding POD were differentially expressed, of which two were significantly downregulated and one upregulated. Of 210 total unigenes encoding serine proteases, 38 were differentially expressed; seven genes belonged to chymotrypsin, 15 to serine protease, and 16 to trypsin. Five chymotrypsin DEGs were upregulated in Sample A, one of which was also upregulated in Sample B; the other two were downregulated in Sample A. All chymotrypsin genes in Sample C and most in Sample B were not significantly differentially expressed (Table 6). Only three genes encoding serine protease were significantly downregulated in all three treatments; the remaining 12 DEGs were variously upregulated, especially in Sample A. Five genes encoding trypsin were upregulated in Sample A, and eight in Sample B; the remaining DEGs were significantly downregulated in Sample A.

**Table 6.**
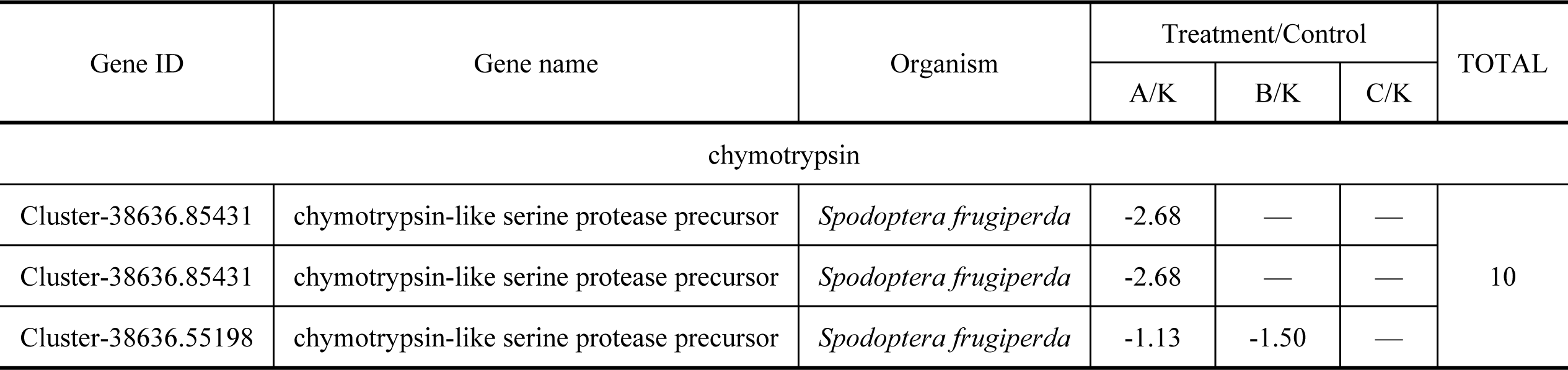
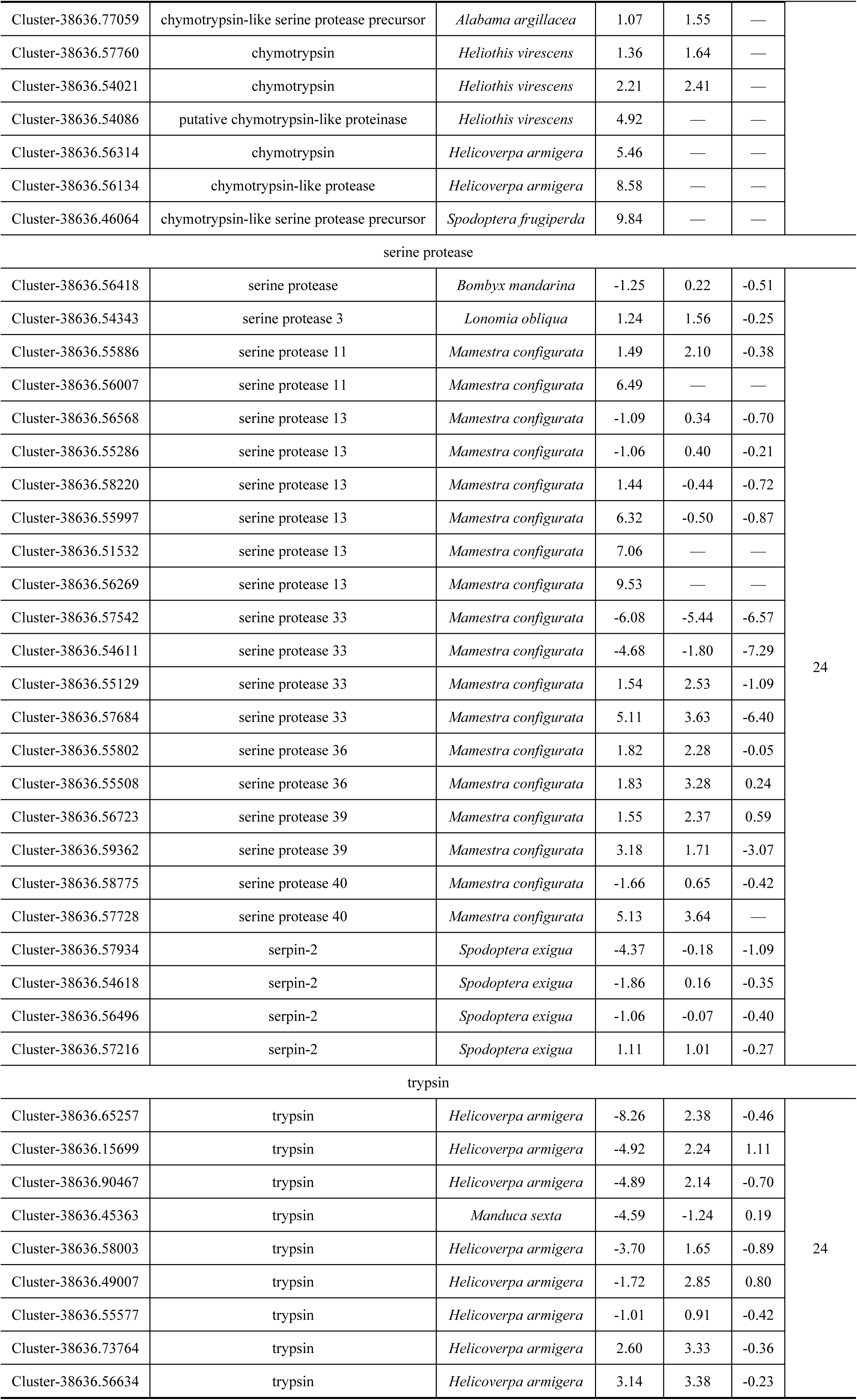
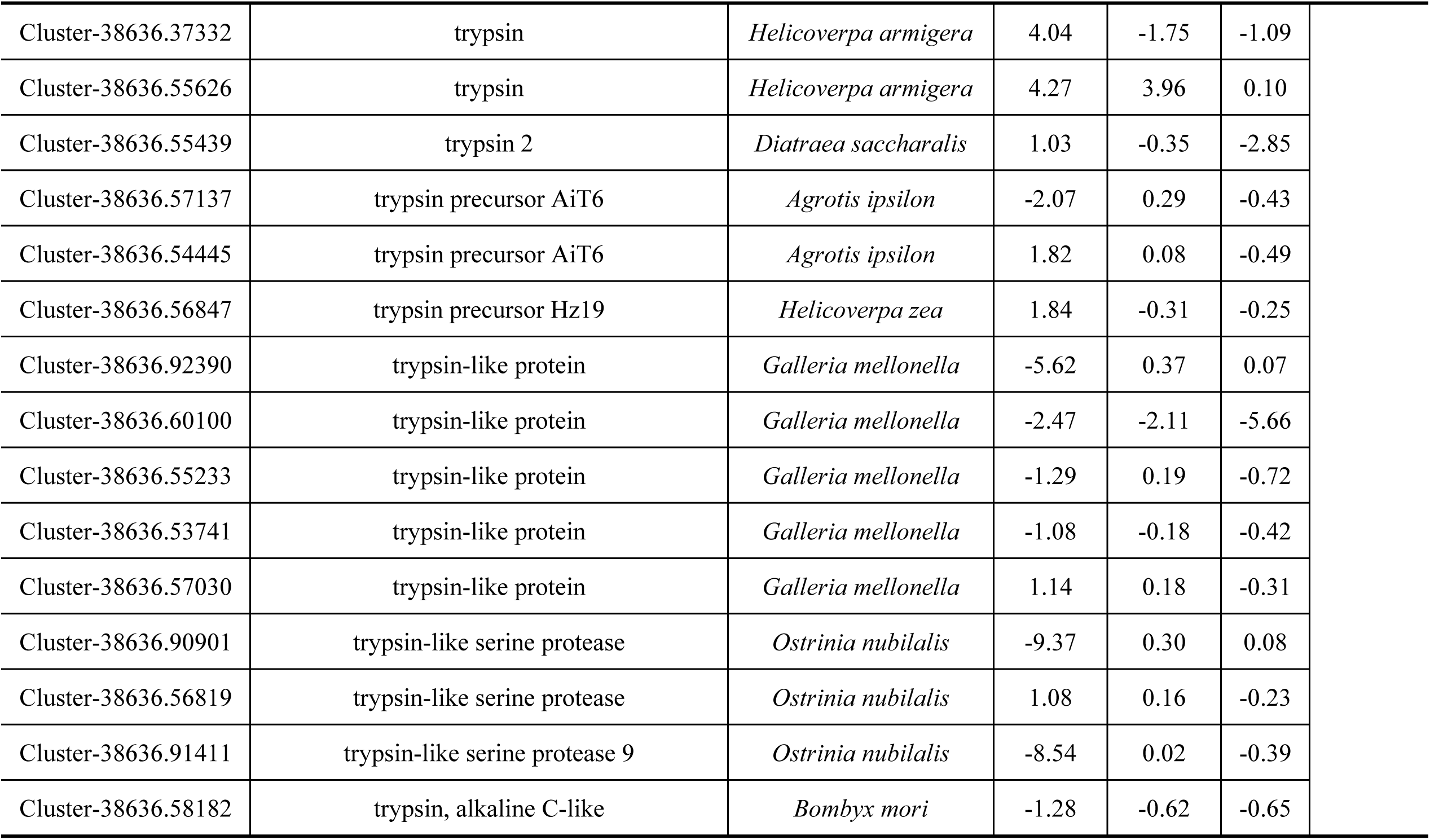
Trypsin genes of the *C. anachoreta* midgut in response to ingestion of Cry1Ac toxin

### 3.9 Pathways significantly enriched in DEGs

Metabolic pathway enrichment analysis revealed 158 common DEGs involved in 102 pathways (Supplementary Table 7) potentially involved in digestion, absorption, and immunity. DEGs were significantly enriched in ten functional categories; six categories, Fructose and mannose metabolism, Glycerolipid metabolism, Glycolysis/gluconeogenesis, Carbohydrate digestion and absorption, Pentose and glucuronate interconversions, and Pyruvate metabolism, are involved in digestion and absorption of food and energy metabolism; two categories, Viral myocarditis and Cardiac muscle contraction, are related to cardiac muscle and may affect blood circulation. The two DEGs involved in the Histidine metabolism pathway might influence production of diapause hormone B, thereby affecting insect development duration. Finally, the Tight junction pathway, a Cell-junction pathway, is common in monolayer columnar epithelium and monolayer cubic epithelium in the midgut, and plays an important role in closing the gap at the top of the cell and blocking entry of extraneous macromolecules into the tissue. All DEGs involved in these pathways might by affected by Cry1Ac toxin (P < 0.05) (Table 8), suggesting that greater energy is needed to sustain life activities when organisms are threatened by drugs or other stressors.

**Table 8.** Pathways enriched in DEGs in *C. anachoreta* larvae treated with Cry1Ac

| Pathway ID | Pathway name | Input number | Background number | Background number | P-value |
| --- | --- | --- | --- | --- | --- |
| ko00051 | Fructose and mannose metabolism | 4 | 158 | 151 | 0.001 |
| ko05416 | Viral myocarditis | 3 | 158 | 127 | 0.0058 |
| ko00561 | Glycerolipid metabolism | 4 | 158 | 290 | 0.0094 |
| ko00010 | Glycolysis / Gluconeogenesis | 4 | 158 | 327 | 0.014 |
| ko04530 | Tight junction | 4 | 158 | 362 | 0.0195 |
| ko04973 | Carbohydrate digestion and absorption | 3 | 158 | 210 | 0.0219 |
| ko00040 | Pentose and glucuronate interconversions | 3 | 158 | 212 | 0.0225 |
| ko00340 | Histidine metabolism | 2 | 158 | 93 | 0.029 |
| ko04260 | Cardiac muscle contraction | 3 | 158 | 236 | 0.0294 |
| ko00620 | Pyruvate metabolism | 3 | 158 | 254 | 0.354 |

### 3.10 Pathways significantly enriched in genes of interest and Bt-related genes

Metabolic pathway enrichment analysis revealed 114 genes of interest involved in 23 pathways (Supplementary Table 9) with important roles in the response of *C. anachoreta* to Cry1Ac toxin. These genes were significantly enriched in nine functional categories, four of which, Glutathione metabolism, Drug metabolism-cytochrome P450, Metabolism of xenobiotics by cytochrome P450, and Chemical carcinogenesis, are related to detoxification and immunity (Table 10).

**Table 10.**
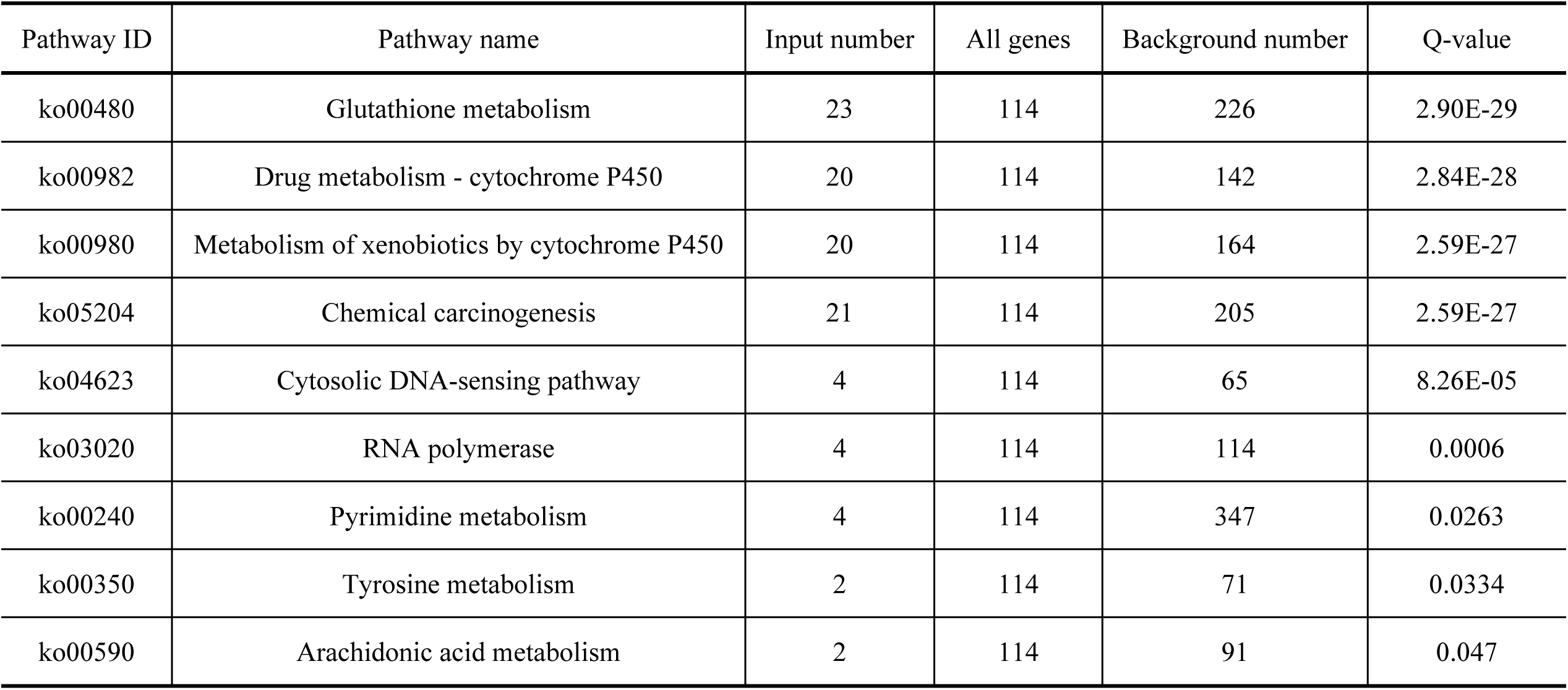
Pathways enriched in genes of interest in *C. anachoreta* larvae treated with Cry1Ac toxin (Q-value < 0.05)

## 4 Discussion

*C. anachoreta* is an important insect pest in forestry, especially in poplar woods in China (except in Xinjiang, Guizhou, and Taiwan), Europe, Japan, and India; it is also the target insect of Cry1Ac toxin and Bt plants. However, the interaction between *C. anachoreta* and Cry1Ac toxin is unclear. The reported half-maximal lethal concentration (LC50) of the DOR Bt-1 formulation was 247.52 µg/mL for 7-day-old or third instar *A. janata* larvae (Vimala Devi and Sudhakar, 2006), and one tenth of the LC50 was used as a sub-lethal dose (24.75 µg/mL) for *A. janata*(Chauhan VK, et al. 2017). In the present study, a 7.5 µg/mL dosage of Bt Cry1Ac toxin solution with sodium carbonate represented a sub-lethal dose, much lower than for other species, suggesting that the *C. anachoreta* larval population used in our experiment was more susceptible to Bt, making it an appropriate choice for studying the mechanism of the response to Bt Cry1Ac.

Possible resistance mechanisms in Lepidoptera include (1) solubilization and incomplete toxin processing and (2) modified Cry toxin binding sites and differential gene expression (Ferré,J. and Van Rie, 2002). The most reported mechanism of resistance seems to be altered binding of Cry toxins to receptors. Notably, a single species can evolve a repertoire of resistance mechanisms to the same or different Cry toxins, such as *H. armigera* 5-405 (NA405), with resistance to Cry2Ab and cross-resistance to Cry2Aa and Cry2Ae; *P. xylostella* (Cry1Ac-R), with resistance to Cry1Ac and cross-resistance to Cry1Ab; and *P. xylostella* NIL-R (BC6F4), with resistance to Cry1Ac and cross-resistance to Cry1Ab and Cry1Ah (B.Peterson et al., 2017).

Based on the transcriptome data, the response of *C. anachoreta* to Cry1Ac toxin is related to its tolerance/resistance and immunity to Bt toxin. In this study, the larval midgut transcriptome was sequenced and analyzed by Novogene, and 151,090 high-quality unigenes were obtained, including 1660, 1175, and 562 DEGs between the three larvae treatments with Cry1Ac toxin and the control, respectively. In the *P. xylostella* midgut transcriptome, 2925 and 2967 unigenes were differently expressed between two resistant strains compared to susceptible strains, respectively(Lei Y, 2014). The DEGs were primarily associated with Bt receptors (e.g., ABC transporter, APN, ALP, and CAD), digestive function (α-amylase, lipase, protease, and carboxypeptidase), immune response (HSC70 and IL22), detoxication enzymes (P450, GST, and CarE), developmental genes (arylphorin), and a variety of binding proteins including cellular retinoic acid-binding protein (lipid-binding), insulin-related peptide-binding protein (protein-binding), and ovary C/EBPg transcription factor (nucleic acid-binding) (Michael E. Sparks, et al,. 2013; Yang Y, et al,. 2018; Juan Luis Jurat-Fuentes, et al,. 2011; ShihoTanaka, et al,. 2013; Josue Ocelot, et al,. 2017; Tristan Stevens et al,. 2017, Jian xiuYao, et al,. 2017; Anais Castagnola et al., 2017; Vinod K. Chauhan et al., 2017). In this study, the common DEGs were also primarily associated with digestion function, with enriched GO terms including monocarboxylic acid metabolic process, pyruvate metabolic process, lipid biosynthetic process, and serine-type endopeptidase activity. We also focused on genes of interest involved in insecticides and Bt resistance or tolerance. These findings may facilitate the understanding of interactions between insects and Bt toxin.

### 4.1 DEGs

There were many DEGs related to growth, development, and metabolism in the midgut of *C. anachoreta.* HSC70, GNB2L/RACK1, and PNLIP were significantly downregulated compared to the control. HSC70 is an important stress-resistance protein against environmental stresses; it is synthesized constitutively in insects and induced by stressors such as heat, cold, crowding, and anoxia.

Several reports demonstrated that HSC70 expression decreased significantly under chlorpyrifos treatment (P < 0.05) (Sun Y et al, 2016). Under exposure to two sub-lethal doses (LC10 and LC30) of β-cypermethrin, both Hsp70-1 and Hsp70-2 expressions reached a maximum 24 h after exposure (Yuting Lia, 2017). HSC70 genes might be highly expressed during the early response to stress, followed by decreased expression during recovery (Allison M, 2015). In our study, HSC70 was significantly downregulated in samples treated with Cry1Ac toxin. HSC70 was involved in six pathways; we focused on the MAPK signaling pathway (ko04010), as it likely plays an important role during Bt metabolism, affecting larval growth and development. GNB2L/RACK1, known as guanine nucleotide-binding protein G (I)/G(S)/G(T) subunit beta-1, binds to and stabilizes activated PKC, increasing PKC-mediated phosphorylation. It recruits activated PKC to the ribosome, leading to phosphorylation of EIF6 and inhibiting the activity of SRC kinases, including SRC, LCK, and YES1. GNB2L participates in the crucial “Chemokine signaling pathway,” which is related to the *C. anachoreta* immune system; downregulation of GNB2L in larvae can result in hypoimmunity, viral infection, or even death. PNLIP encodes a lipase family protein; lipases are essential for the efficient digestion of dietary fats. Downregulation of PNLIP might result in metabolic disorders, subsequently affecting larval growth and development.

The unigenes encoding Bax inhibitor-1-like protein (BI-1), arylphorin type 2, and pyruvate kinase (PKM) were significantly upregulated in midgut treated with Cry toxin. BI-1 is one of the few cell death suppressors conserved in animals and plants, with a membrane-spanning protein with six to seven transmembrane domains and a cytoplasmic C-terminus within the endoplasmic reticulum and nuclear envelope. BI-1 is involved in development and response to biotic and abiotic stress, and probably represents an indispensable cell protectant that appears to suppress cell death induced by mitochondrial dysfunction, reactive oxygen species, or elevated cytosolic Ca^2+^ levels (Hückelhoven R, 2004). Significant upregulation of BI-1 might prevent larval cell death due to poisoning from Cry toxin, imposing a certain fitness cost after survival, such as loss of weight, lengthened development period, etc. Alpha-arylphorin is a mitogen in the *H. virescens* midgut cells secretome upon Cry1Ac intoxication(AnaisCastagnola, 2017), and the most significant difference detected in the Cry1Ac secretome was an arylphorin subunit alpha protein not detected in the control secretome, which caused higher proliferation and differentiation in primary midget stem cells. In the cases of *S. exigua* and *Diatraea saccharalis*, arylphorin genes were upregulated in response to intoxication with Bt toxin (Hernández-Martínez et al., 2010; Guo et al., 2012). Previous reports were consistent with our results showing that arylphorin type 2 and arylphorin subunit beta-like were significantly upregulated in treatments compared to the control, contributing to the molecular regulation of midgut healing in the larval response to Cry toxin. The final significantly upregulated DEG encoded PKM, which is involved in the fifth step of the glycolysis sub-pathway that synthesizes pyruvate from D-glyceraldehyde 3-phosphate, a part of the carbohydrate degradation pathway. In our study, PKM participated in six pathways, four of which (Glycolysis, Pyruvate metabolism, Carbon metabolism, and Biosynthesis of amino acids) are related to carbohydrate degradation and energy metabolism, suggesting that larvae require more energy to stay alive when suffering from stress.

In summary, whether upregulated or downregulated, the DEGs participated a variety of pathways, such as the immune pathway, detoxification metabolism, etc., to regulate cells or life processes, allowing the larvae to remain alive. However, the data are insufficient to determine the specific mechanisms involved in pathways that play crucial roles in the midgut of *C. anachoreta* after treatment with Cry toxin.

### 4.2 DEGs in pathways of interest

The mechanism of action of Cry toxins has been studied for many years (Pigott and Ellar, 2007; Whalon and Wingerd, 2003; Pardo-Lopez et al., 2013). Multiple studies have demonstrated that the MAPK p38 pathway is activated in multiple organisms/insect orders (Lepidoptera, Coleoptera, and Diptera) in response to a variety of pore-forming toxins, including Cry1A. This pathway activates a complex defense response (Leivi P. et al. 2016; GuoZ et al. 2015; Angeles Cancino-Rodezno et al.2010), including interaction with cadherin receptor, triggering an intracellular cascade response involving protein G, adenylate cyclase, and protein kinase A. In *Caenorhabditis elegans* both the MAPK p38 and c-Jun N-terminal kinase (JNK) pathways have been reported as important to the defense response against Cry5B toxin (Huffman et al., 2004). In *B. mori*, the JNK and JAK-STAT pathways were found to be upregulated in the early response to Cry1Aa intoxication (Tanaka, Yoshizawa &Sato, 2012).

Here, the response to Bt Cry1Ac toxin might involve the PI3K-Akt pathway, which was highly enriched in DEGs and linked to several crucial pathways, including the B cell receptor signaling pathway, toll-like receptor pathway, and MAPK signaling pathway. The B cell receptor and toll-like receptor pathways are related to adaptive and innate immunity in the early response to Cry toxin, while the MAPK signaling pathway is involved in the mechanism of resistance in resistant insects. In our study, the PI3K-Akt pathway was activated by a FAK tyrosine kinase (also known as RTK), which was stimulated via the recognition of integrin beta-1 and Von Willebrand factor; the genes of all three were upregulated. FAK is required during development and might regulate tumor suppressor p53. The binding of growth factors to RTK stimulates class Ia PI3K isoform, and PI3K catalyzes the production of phosphatidylinositol-3,4,5-triphosphate (PIP3) at the cell membrane. PIP3 in turn serves as a second messenger that helps to activate Akt. Once active, Akt can control key cellular processes by phosphorylating substrates involved in apoptosis, protein synthesis, metabolism, and cell cycling. In our study, the downregulated AKT gene might have encoded less activated Akt, thereby reducing phosphorylation of downstream factors, and increasing downstream reactions such as BAD or Casp9 activation of the apoptosis process. It could also result in the accumulation of PEPCK, indirectly activating the process of glycolysis, which could generate more ATP (energy) and NADH (reducing power) for other life activities. Our results differed from those from other resistant insects, possibly due to the timing of midgut dissection, which took place after a period of recovery. During the recovery period until growth to fourth instar, the larvae dieted on control leaves. However, due to damage to the midgut caused by the pore-forming toxin, the larvae required more energy to digest and absorb nutrition from food, and to heal midgut epithelial cells through proliferation and apoptosis.

Therefore, in the midgut of *C. anachoreta* larvae treated with Bt Cry1Ac and recovered to fourth instar, the glycolysis and oxidative phosphorylation pathways might be activated to generate more ATP (energy) and NADH (reducing power) for larval growth and development. Meanwhile, apoptosis and regeneration pathways might be stimulated and upregulated for recovery of the larval midgut.

